## Supplementary Material for "Virus-host dynamics in archaeal groundwater biofilms and the associated bacterial community composition"

Content:

1. Supplementary Tables
2. Supplementary Figures
3. Supplementary References

Supplementary Tables are provided as Supplementary_Tables.xlsx file, which contains the following individual sheets:

- **Table S1:** Sampling events on the Mühlbacher Schwefelquelle (MSI).
- **Table S2:** Melting profile of the primer set Altivir_1_MSI_F and Altivir_1_MSI_R for targeting the lytic virus Altivir_1_MSI in individual biofilm flocks from the MSI.
- **Table S3:** qPCR standard curves of the respective primer sets for targeting *Ca.* Altiarchaeum hamiconexum, Altivir_1_MSI and bacteria/archaea in individual MSI biofilm flocks
- **Table S4:** Direct-geneFISH probes for targeting the genome of *Ca.* A. hamiconexum for the determination of the detection efficiency of virusFISH.
- **Table S5:** Melting profiles of the direct-geneFISH probes for targeting the genome of *Ca.* A. hamiconexum for the determination of the detection efficiency of virusFISH
- **Table S6:** Determining the detection efficiency of virusFISH (raw data).
- **Table S7:** Calculation of the detection efficiency of virusFISH (results are displayed in Fig. 2).
- **Table S8:** Kruskal-Wallis and Dunn's significance tests for qPCR and virusFISH data sets.
- **Table S9:** Nanopore sequencing - Barcode sequences for the16S Barcoding Kit.
- **Table S10:** Raw data of virus-host ratios of different methods, real-time PCR, metagenomics, and virusFISH (results are illustrated in Fig. 1B).
- **Table S11:** Viral enumeration of different infection stages with Altivir_1_MSI (results are visualized in Figure 1A).
- **Table S12:** Targeting *Ca.* A. hamiconexum, its virus Altivir_1_MSI and the entire bacteriome within individual MSI biofilms by using real-time PCR (results are illustrated in Fig. 3).

**Figure S1:** VirusFISH on MSI biofilms by using a non-matching *Metallosphaera* sp. virus probe as a negative control for **Main Figure 3**.

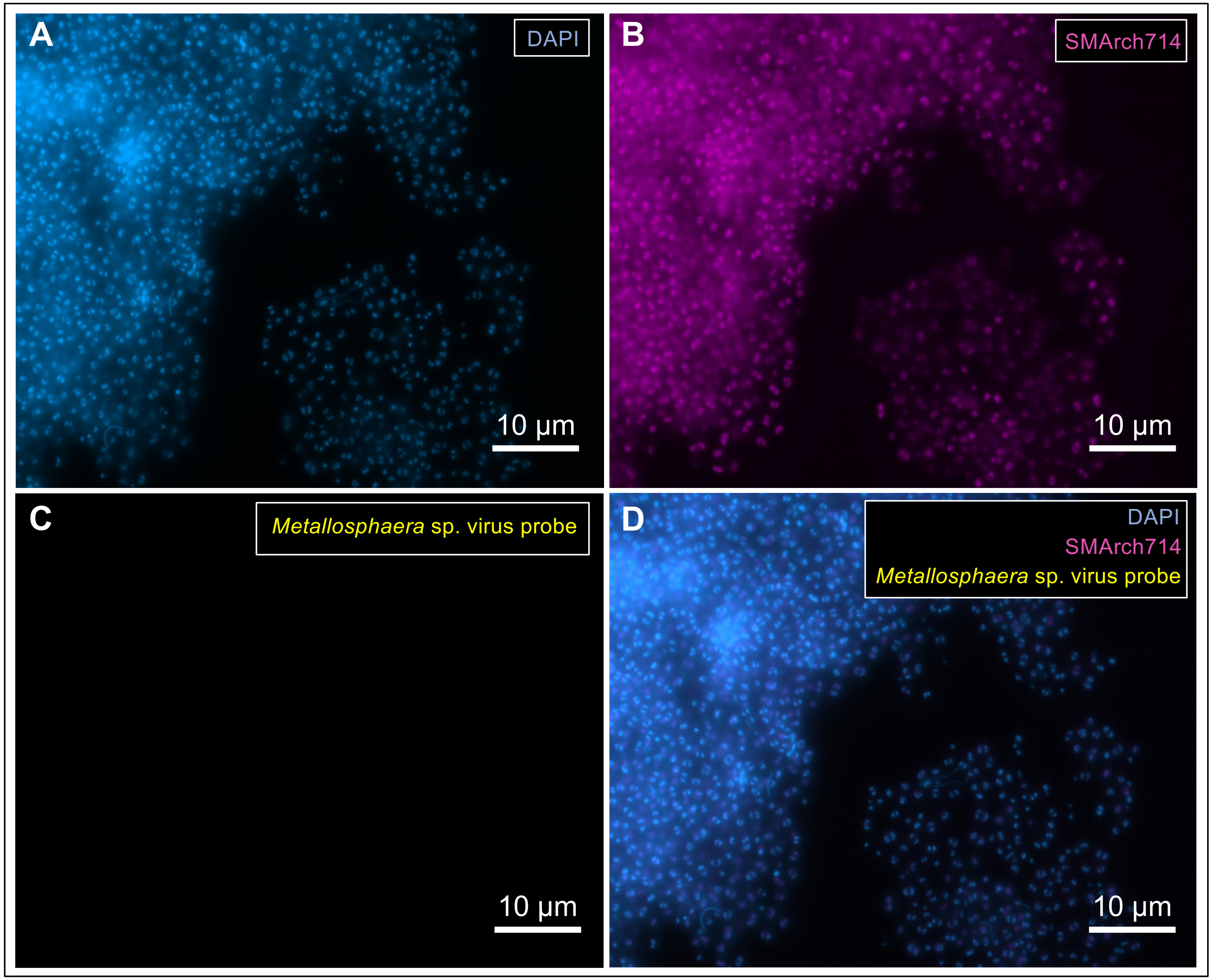

**Figure S2:** Direct-geneFISH of *E. coli* cells as a negative control for **Main Figure 2C** with all 33 different polynucleotides that specifically target the *Ca.* Altiarchaeum genome.

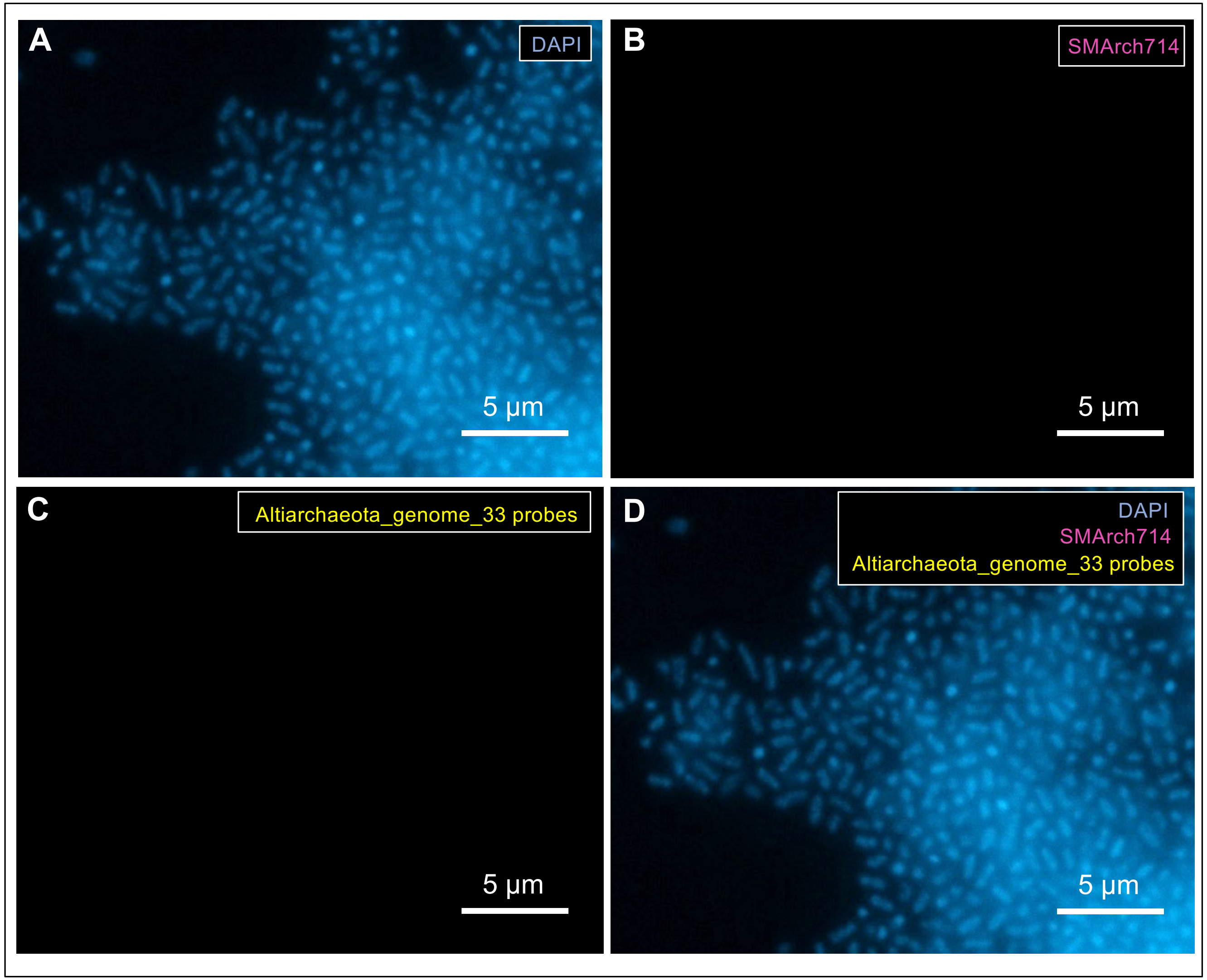

**Figure S3:** Extended data of **Main Figure 2C**, methods according to main manuscript. Direct-geneFISH on MSI biofilms using eleven different polynucleotides to target the *Ca.* Altiarchaeum genome. Only strong, punctual signals were counted for calculating the labelling efficiency.

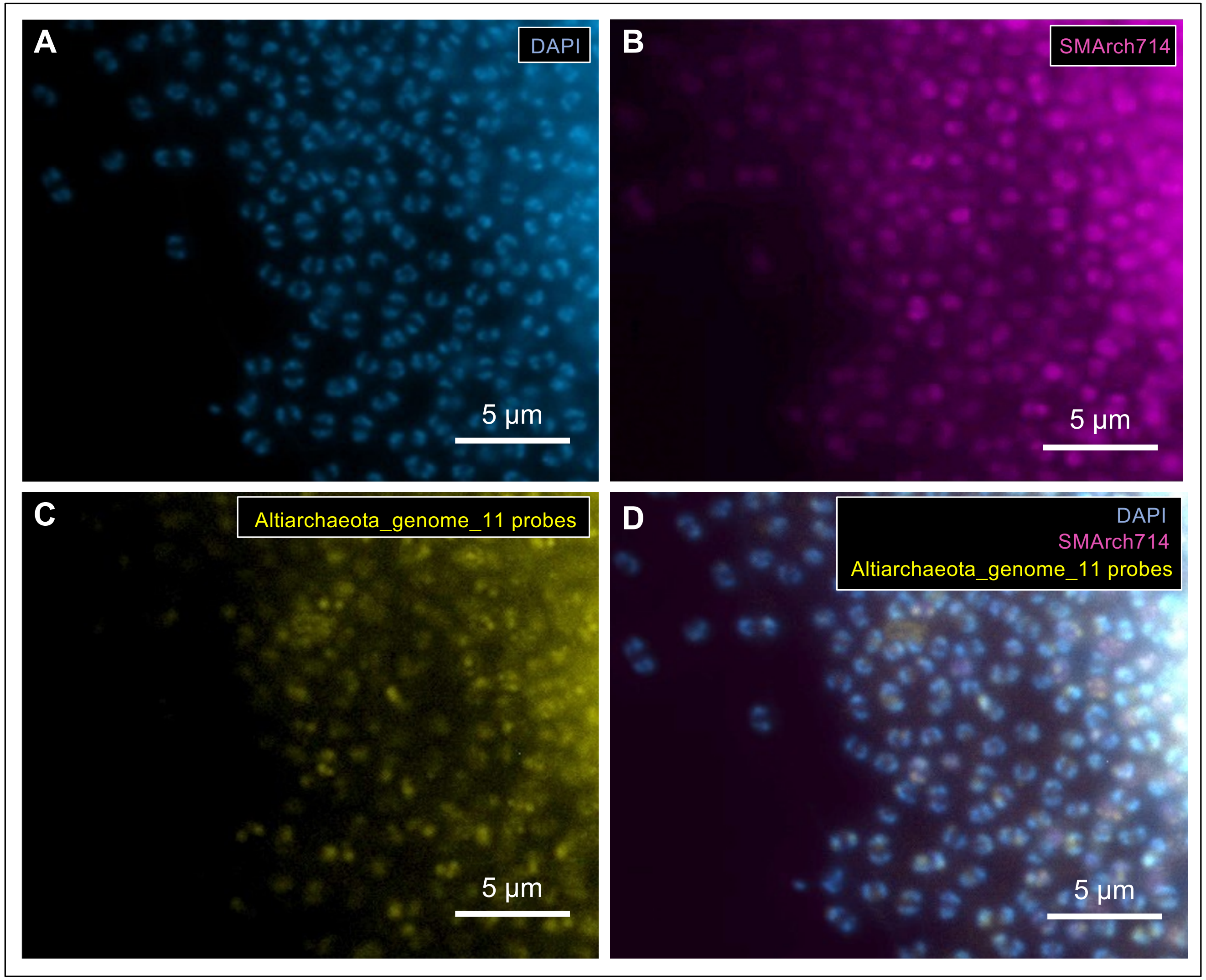

**Figure S4:** Extended data of **Main Figure 2C**, methods according to main manuscript. Direct-geneFISH on MSI biofilms using 22 different polynucleotides to target the *Ca.* Altiarchaeum genome. Only strong, punctual signals were counted for calculating the labelling efficiency.

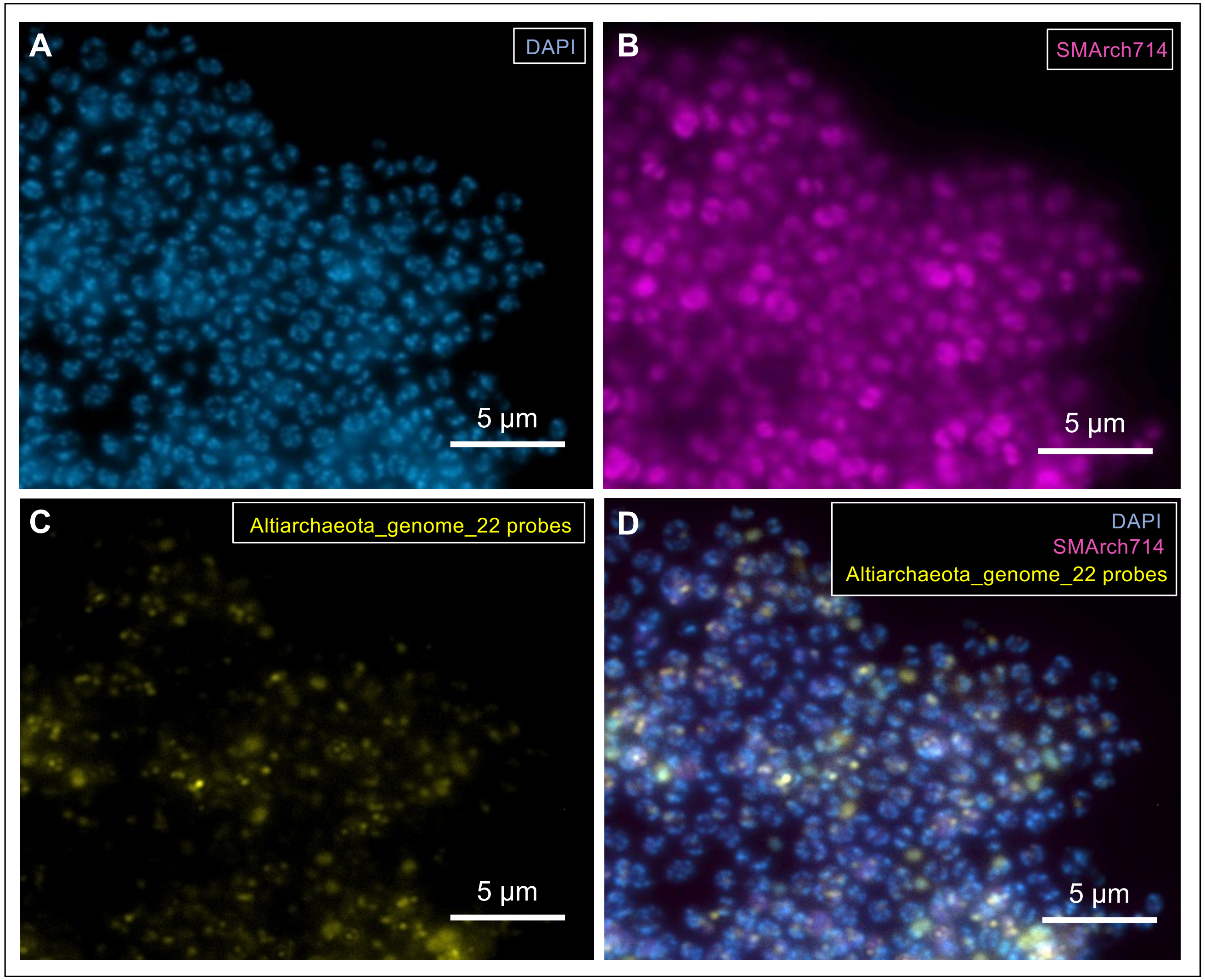

**Figure S5:** Extended data of **Main Figure 2C**, methods according to main manuscript. Direct-geneFISH on MSI biofilms using 33 different polynucleotides to target the *Ca.* Altiarchaeum genome. Only strong, punctual signals were counted for calculating the labelling efficiency.

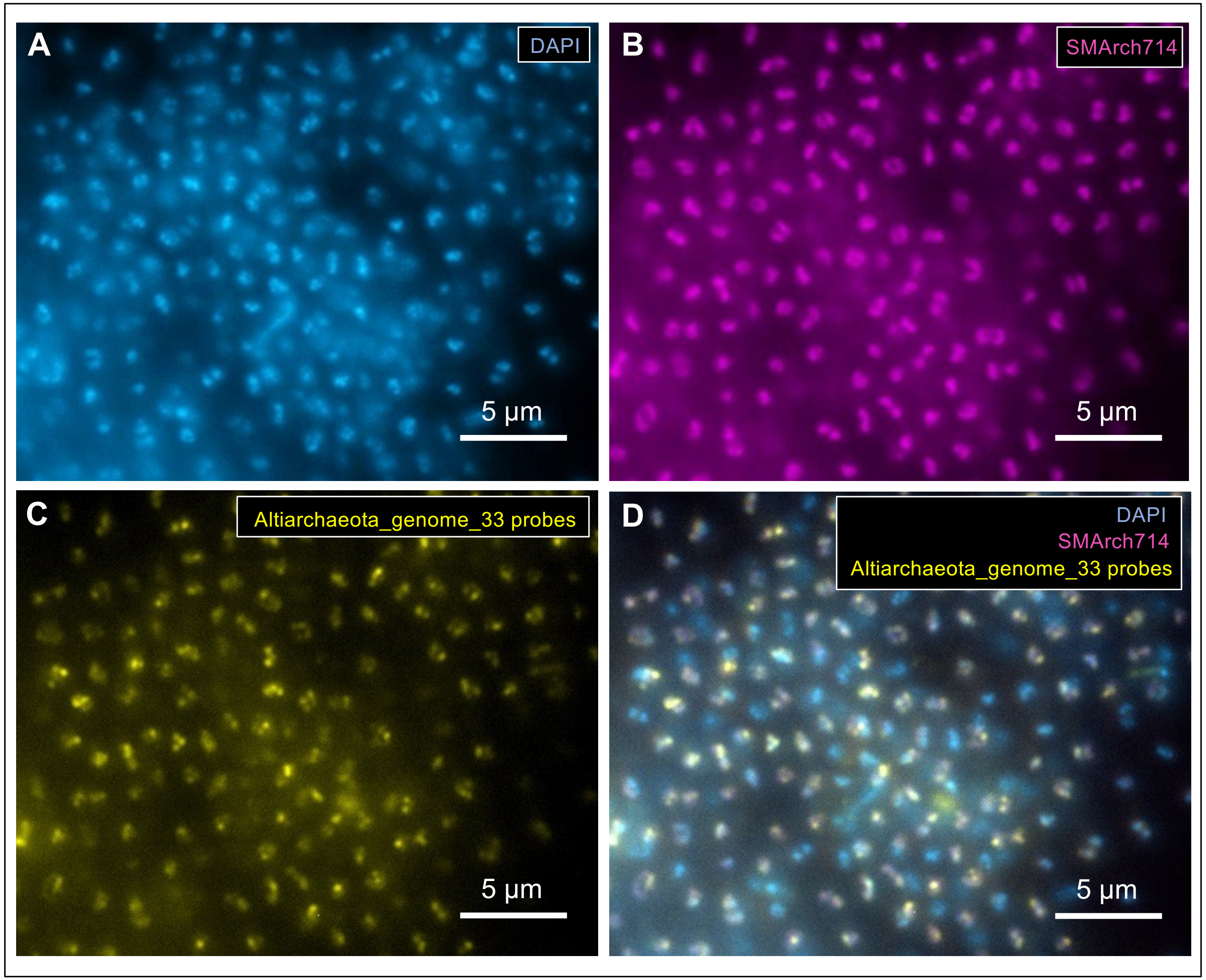

**Figure S6:** Extended data of **Main Figure 3A**, methods according to main manuscript. VirusFISH on MSI biofilms shows few viral infections caused by Altivir_1_MSI.

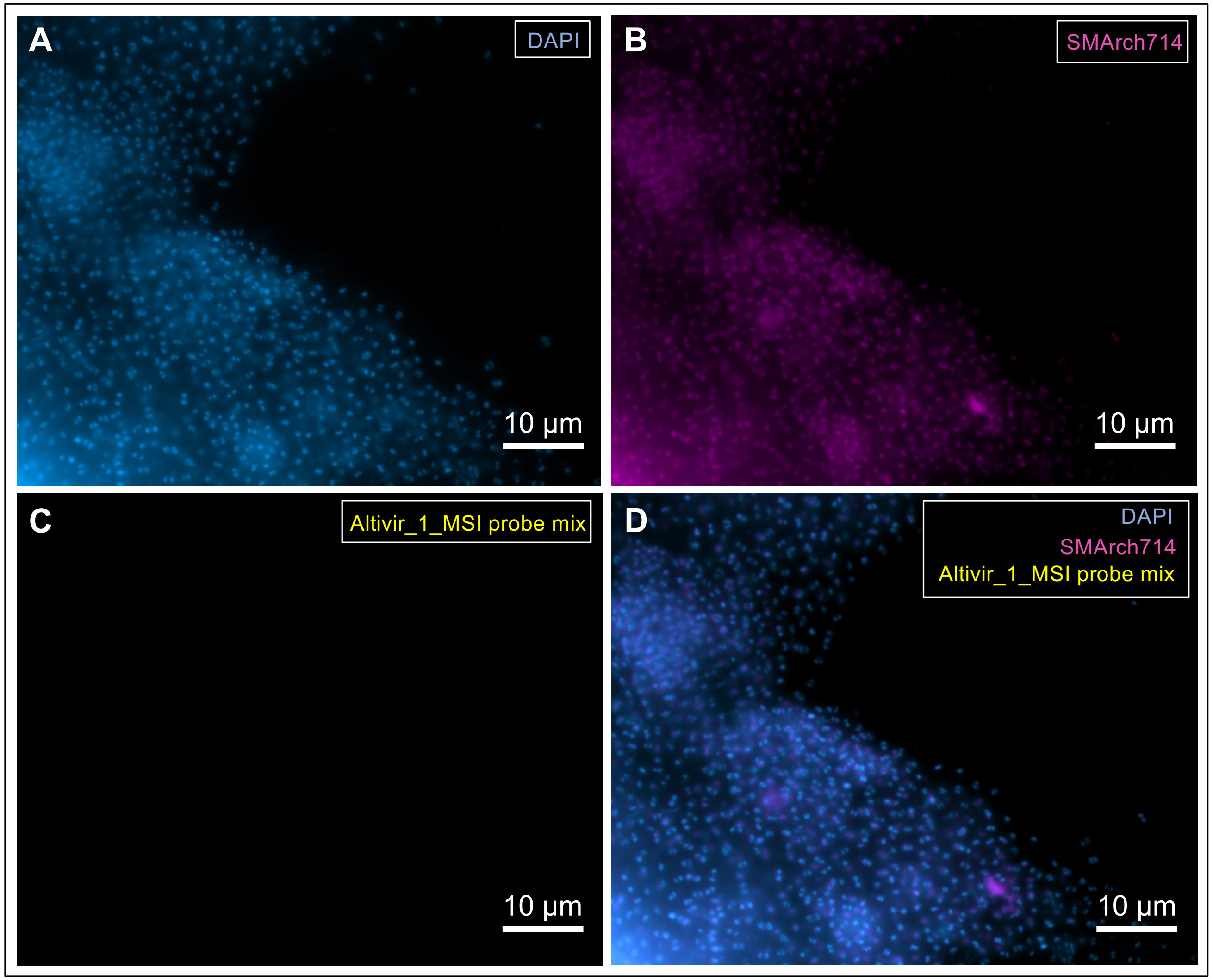

**Figure S7:** Extended data of **Main Figure 3B**, methods according to main manuscript. VirusFISH on MSI biofilms shows an increase in the infection frequency caused by Altivir_1_MSI.

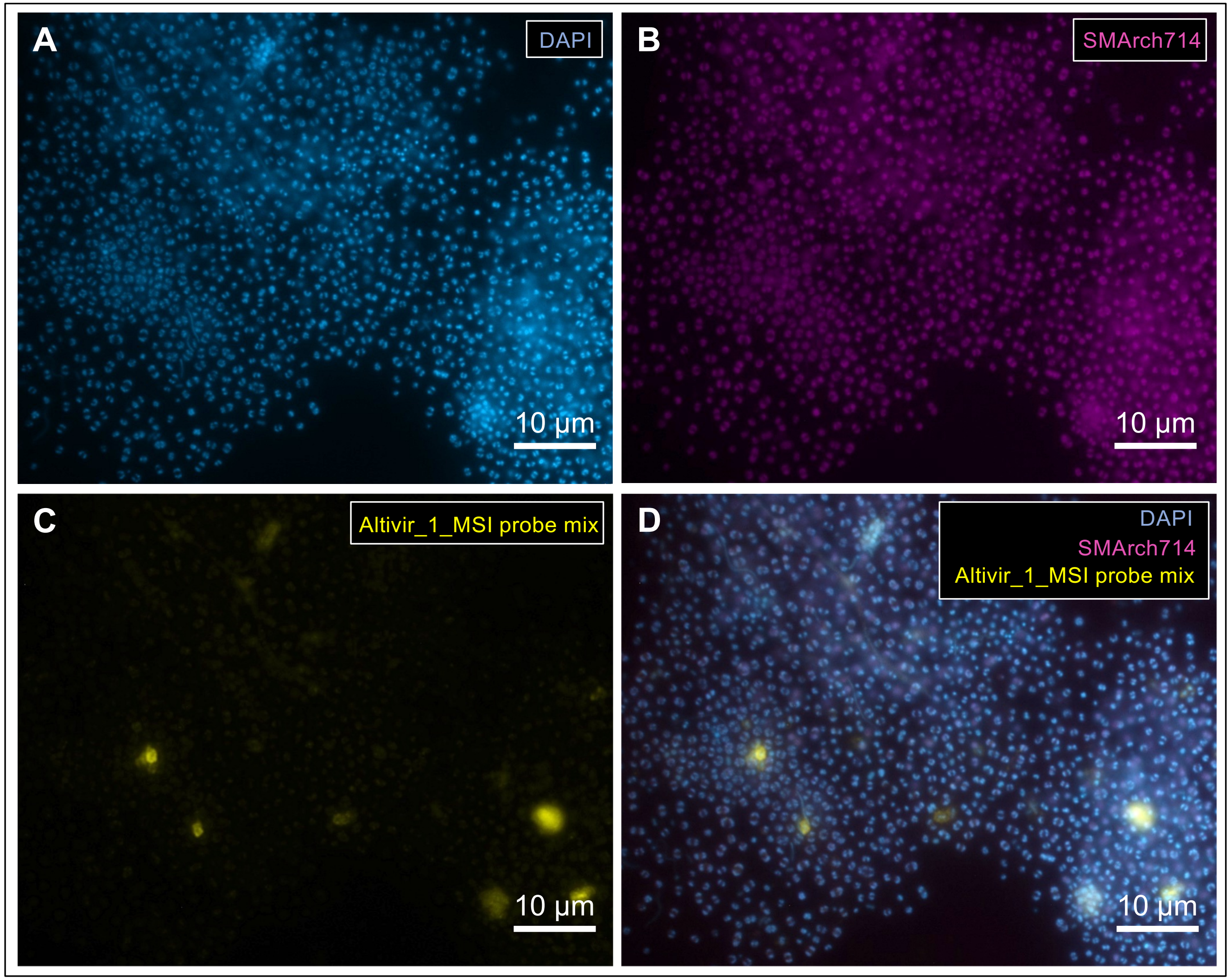

**Figure S8:** Extended data of **Main Figure 3C**, methods according to main manuscript. VirusFISH on MSI biofilms shows that the vast majority of the host cells are infected by Altivir_1_MSI.

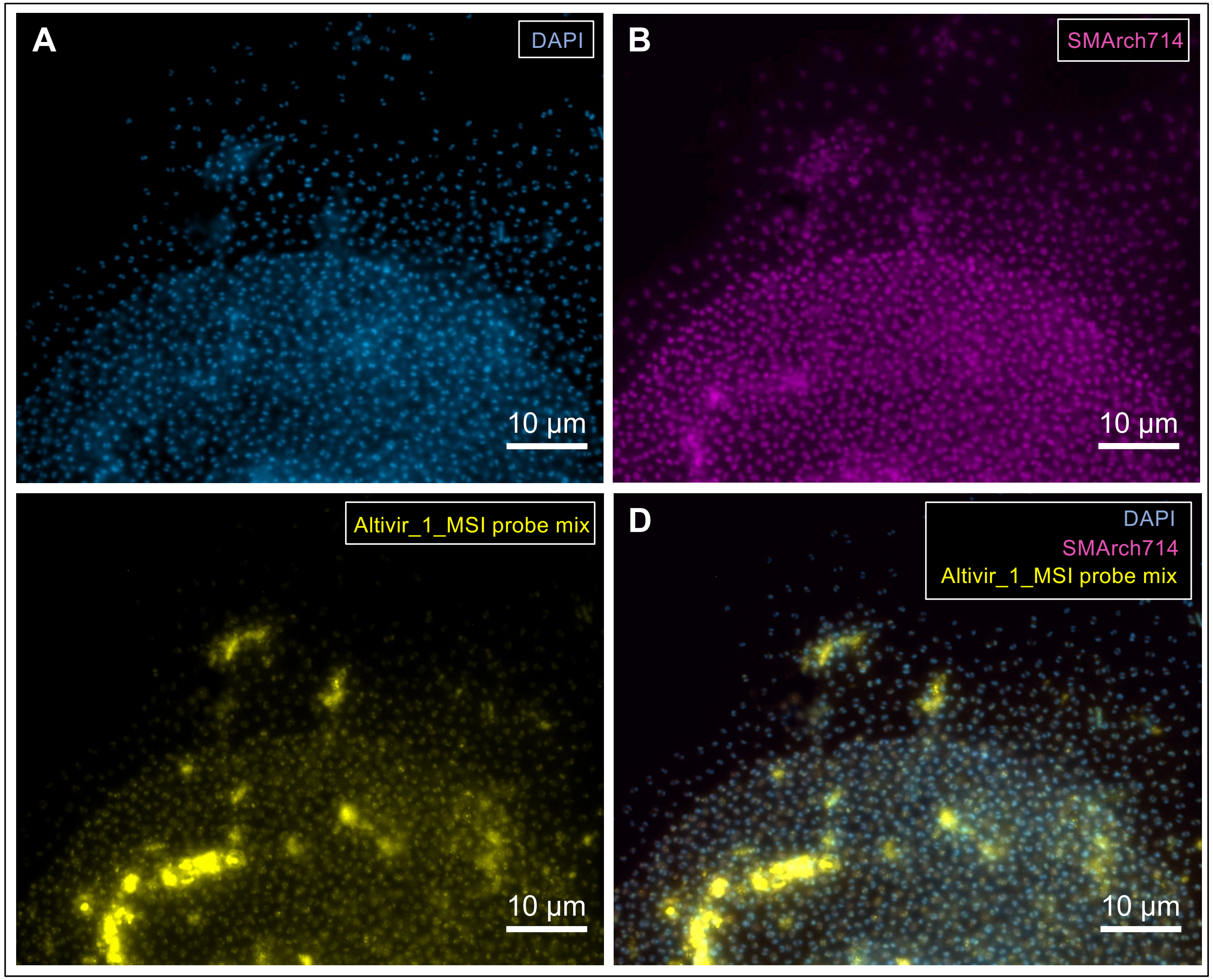

**Figure S9:** Extended data of **Main Figure 3D**, methods according to main manuscript. VirusFISH on MSI biofilms shows cell lysis caused by Altivir_1_MSI and the enrichment of filamentous microbes along with the cell debris.

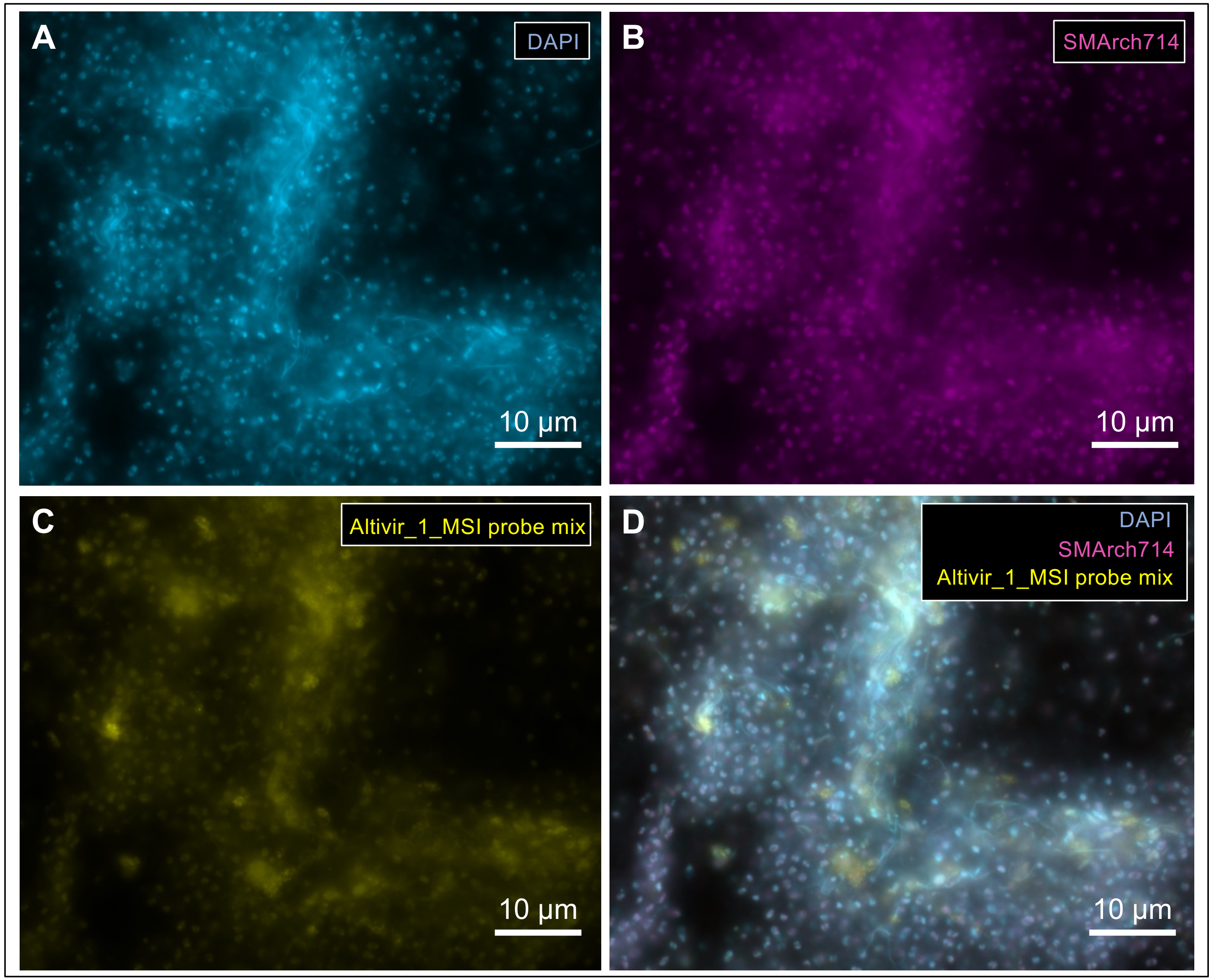

**Figure S10:** Extended data of **Main Figure 4**. Correlation of the 16S rRNA gene relative abundances with the virus-host ratio showing no linear relationship among the most abundant bacterial taxa in individual MSI biofilm flocks.

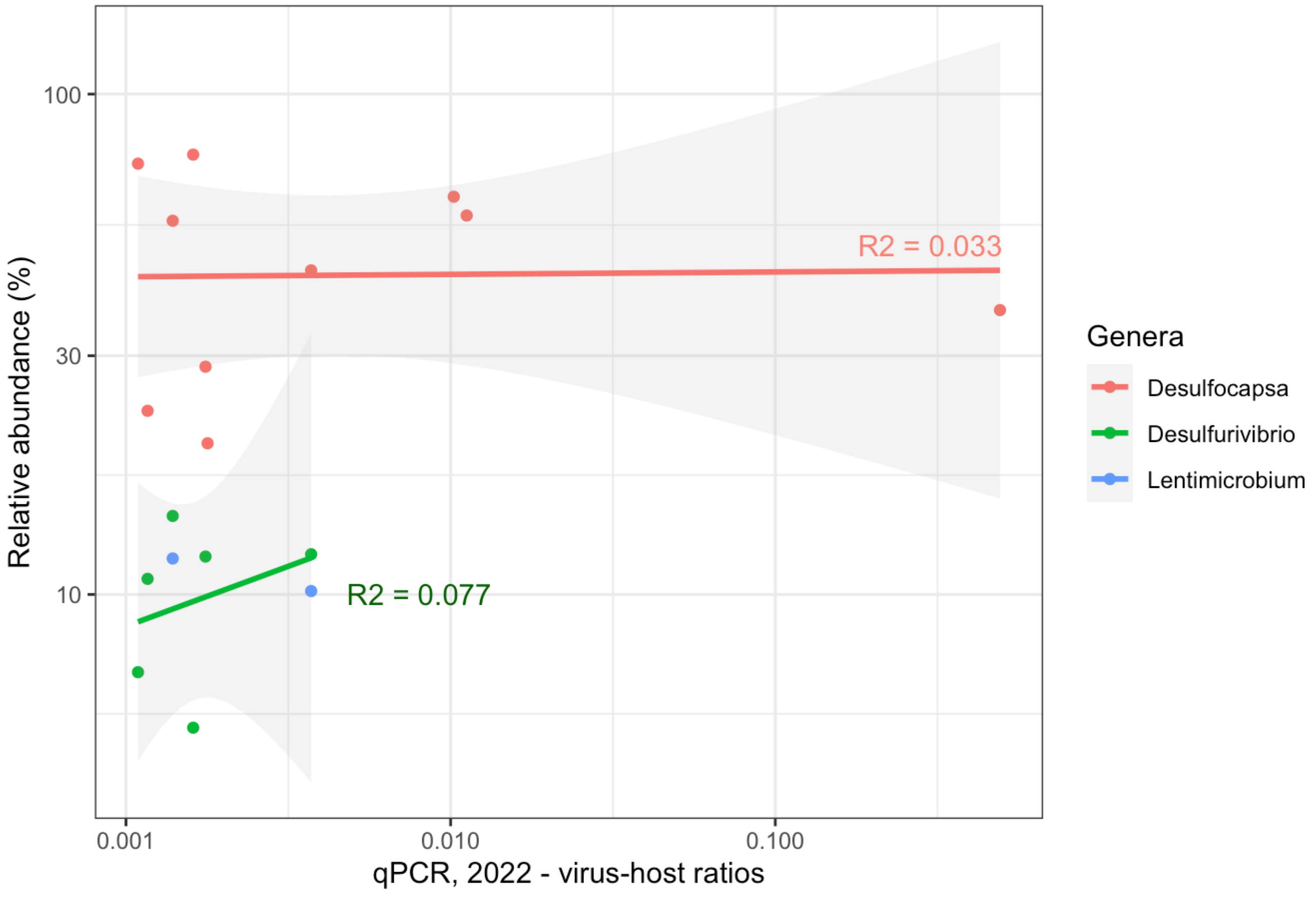
